# Baseline dependent differences in the perception of changes in visuomotor delay

**DOI:** 10.1101/2024.06.20.599936

**Authors:** Gesche Vigh, Jakub Limanowski

## Abstract

The detection of, and adaptation to delayed visual movement feedback has been extensively studied. One important open question is whether the Weber-Fechner Laws hold in the domain of visuomotor delay; i.e., whether the perception of changes in visuomotor delay depends on the amount of delay already present during movement. To address this, we developed a virtual reality based, continuous hand movement task, during which participants had to detect changes in visuomotor mapping (delay): Participants (N=40) performed continuous, auditory-paced grasping movements, which were measured with a data glove and transmitted to a virtual hand model. The movements of the virtual hand were delayed between 0-700ms with the delay changing repeatedly in a roving oddball design. Participants had to indicate any perceived delay changes by key press. This design allowed us to investigate detection accuracy and speed related to the magnitude of the delay change, and to the “baseline” delay present during movement, respectively. As expected, larger delay changes were detected more accurately than smaller ones. Surprisingly, delay changes were detected more accurately and faster when participants moved under large > small delays. These results suggest that visual movement feedback delay indeed affects the detection of changes in visuomotor delay, but not as predicted by the Weber-Fechner Laws. Instead, bodily action under small delays may have entailed a larger tolerance for delay changes due to embodiment-related intersensory conflict attenuation; whereas better change detection at large delays may have resulted from their (visual) saliency due to a strong violation of visuomotor predictions.

## 1. Introduction

For much of our usual sensory input, like sound, light and weight, the perceived magnitude of a change is proportional to the initial stimuli perceived before the change. Thus, the larger an initial stimulus is, the larger the change needs to be for the resulting stimulus to be perceived as different. This relationship is described by the Weber-Fechner Laws (Fechner, 1966), which have been confirmed across different sensory modalities (Baird, 1997; Gescheider, 1997). Recent empirical evidence suggests that the Weber-Fechner Laws may also apply to more abstract domains like time and reward perception (Brannon et al., 2008; Namboodiri et al., 2014; Takahashi, 2005, 2006); Nieder and Miller, (2003).

The Weber-Fechner-Laws have also been evaluated in the domain of bodily perception. Interestingly, studies investigating self-motion perception found that the relationship between stimulus intensity and change perception was better described by power laws than the Weber-Fechner-Laws (Mallery et al., 2010; Nesti et al., 2015; Nouri & Karmali, 2018). Carriot and colleagues (2021) found improved discrimination performance at higher stimulus levels (of self-motion), rather than nonlinearities predicted by the Weber-Fechner-Laws. Furthermore, Ganel and colleagues (2008) showed that just noticeable differences depended on object size only in perceptual tasks, not in visuall guided grasping tasks—concluding that the visual coding for action does not follow the Weber-Fechner-Laws. Thus, it is possible that the processing of stimuli related to bodily action and self-motion may follow different perceptual laws than the perception of external stimuli (Carriot et al., 2021). One key question that, to our knowledge, has not yet been addressed is whether the Weber-Fechner Laws hold for the perception of visuomotor delays.

The flexible control of bodily action by the brain relies on the capacity to adapt internal models for motor control to novel sensory movement feedback (Limanowski, 2022; Shadmehr et al., 2010; Wolpert & Kawato, 1998). Sensorimotor adaptation is necessary in cases where e.g. visual movement feedback is delayed, i.e., a temporal incongruence is introduced between movements and their visual feedback (Foulkes & Miall, 2000; Rohde & Ernst, 2016). Sensorimotor delays result ‘naturally’ from conduction delays within our nervous system, thus posing a problem for the brain, particularly regarding the on-line control of action using (delayed) sensory feedback. It is generally agreed upon that the brain overcomes such delays through internal forward modeling of the sensory movement consequences (Miall & Wolpert, 1996; Todorov & Jordan, 2002). But sensorimotor delays are also increasingly encountered in new technologies enabling cyber-physical interactions, such as virtual reality or robotics (Peters et al., 2018; Schulze et al., 2024). In most of these scenarios, sensorimotor delays are unavoidable (e.g., due to limitations in bandwidth or computing power) and, more importantly, they can vary unpredictably (Planthaber et al., 2018).

Many studies have investigated the perceptual thresholds for detecting visuomotor delays, the capacity to adapt to them, and the neuronal underpinnings of these processes (Farrer et al., 2008; Foulkes & Miall, 2000; Krugwasser et al., 2019; Leube et al., 2003; Limanowski, 2017; Miall et al., 1985; Miall & Jackson, 2006; Parvin et al., 2024; Rohde & Ernst, 2016; Tsakiris et al., 2006). However, to our knowledge, it has not yet been addressed is whether the perceptual sensitivity to visuomotor delay changes depends on the amount of delay present during movement, as predicted by the Weber-Fechner-Laws.

Therefore, we addressed this question through a virtual reality based hand movement task: Participants performed continuous right-hand grasping movements, paced by an auditory rhythm. Their movements were fed to a photorealistic virtual hand model presented on screen, with an experimentally added delay that changed in a roving oddball fashion (Fig. 1). Participants had to report any perceived delay changes (i.e., changes in the mapping of their real hand movements to those executed by the virtual hand) by key presses, as fast as possible throughout the movement task. We expected larger delay changes to be detected more accurately and faster. Furthermore, following the Weber-Fechner Laws, we expected that participants would detect delays better and quicker if they were moving under relatively smaller delays.

**Figure 1.**
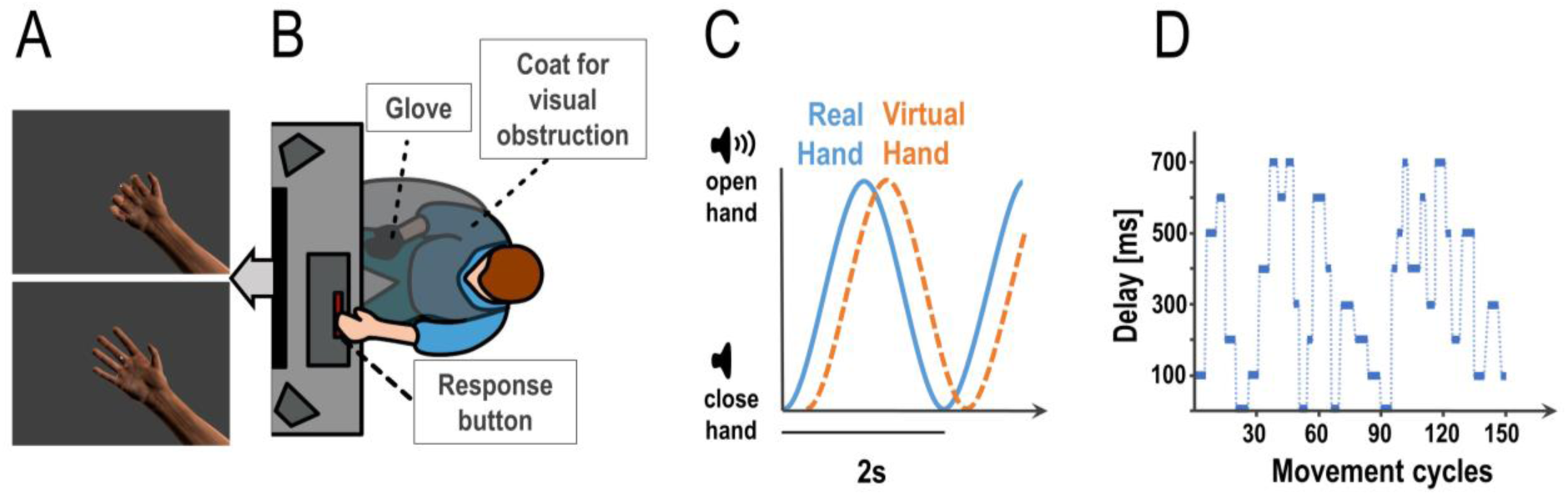
Experimental task and design. Participants controlled the movements of a photorealistic virtual hand presented on screen (**A**) via their hand movements measured by a data glove worn on their right hand (occluded from view, **B**). **C:** Participants had to execute continuous grasping (open-and-close) movements paced by an auditory cue, a tone changing volume with a frequency of 0.5 Hz (i.e., periodically growing louder and quieter). Flexion data from the glove were fed to the virtual hand model, which displayed the executed finger movements. During the task, we added various amounts of delay to the virtual hand movements. This altered the visuomotor mapping; i.e., the temporal congruence between the participants’ executed hand movements and the movements of the virtual hand (i.e., the visual movement feedback). Shown here is a schematic plot showing the real hand closing and opening following the auditory cue’s volume, with a delay added to the virtual hand movements. **D:** During the continuous open-and-close movement task, we repeatedly changed the amount of visual feedback delay in a roving oddball design. This plot shows an example sequence of delays between 0-700 ms as used in the experiment (i.e., one run of 4.9 min = 147 open-and-close movement cycles à 2s each). Participants had to report any perceived changes in the visuomotor delay by pressing a button with their left hand as fast as possible (see **B**).

## 2. Methods

### 2.1 Participants

We recruited 40 healthy, right-handed participants, with normal or corrected-to-normal vision (15 female, mean age = 28.1, range= 22-38). We chose our sample size roughly following previous work investigating sensorimotor conflicts with continuous movement paradigms (Charalampaki et al., 2024; Dewey, 2023; Limanowski & Friston, 2020b; Haering & Kiesel, 2016; Asai, 2015; Salomon et al., 2013). Participants were recruited through undergraduate psychology lectures and seminars and social media. We excluded three participants (one because the data glove could not be adequately fitted and calibrated; one participant was not able to perceive the changes in volume of the tracking sound; and one did not perceive any delay of the virtual hand, even for a delay of 700ms), leaving us with 37 participants whose data could be analyzed. The experiment was approved by the ethics committee of the Technische Universität Dresden and conducted in accordance with this approval.

### 2.2 Experimental Procedure

First, participants were briefed about the experimental procedure and completed a demographic questionnaire. Then they put on a sanitary glove (for hygienic reasons) and the data glove on their right hand (5Dt Data Glove Ultra, 14 sensors, 10 bit flexure resolution per sensor, 60 Hz sampling rate, communication with the PC via USB). The data glove allowed our participants to control a photorealistic virtual hand (Fig. 1B; modelled in blender 2.79, https://www.blender.org/), by measuring the flexion of the participants’ fingers and transmitting these values to the virtual hand model’s fingers (Fig. 1A-C). The virtual hand was shown on a monitor in front of the participant (1920 × 1080 pixels resolution, screen distance to eye about 90 cm, 18.92° of visual angle, 13 ms Motion Picture Response Time). Participants placed their right hand on their leg, with the palm facing upward, in a position that allowed for comfortable opening and closing of the hand (see Fig. 1A). Their right hand was occluded from view by a barber coat. The data glove was then calibrated; i.e., the virtual fingers’ movement range was dynamically adjusted to the sensor values until it plausibly displayed the movements of the real hand.

Participants were instructed to fixate a dot presented centrally on screen, and visible at all times, for the entirety of the experiment (Fig. 1B). The movement task was to execute simple, continuous grasping movements: Participants opened and closed their right hand (starting from a closed position) paced by a rhythmic auditory volume cue; i.e., a 250 Hz tone that grew louder and quieter following a sine wave function with 0.5 Hz frequency. When the tone was loudest, the physical hand of the participant was supposed to be open, when it was quietest the hand was supposed to be closed (Fig. 1C).

Before the actual experiment, participants practiced to align their real hand grasping movement with the auditory rhythm (without added delay to the virtual hand) until they felt confident that they could follow the auditory cue with their hand movements continually throughout the main experiment.

In the first part of the experiment, we tested for potential differences in the perceived difficulty of the hand movement task itself across the different delay levels used in the main experiment. Participants had to perform 10 auditory paced grasping movements (20 s) while fixating the dot in front of the virtual hand, at each delay level used in the main experiment – and rate the task difficulty (see below). I.e., the virtual hand moved either synchronously (0 ms delay) or was delayed by 100, 200, 300, 400, 500, 600, or 700 ms. We chose relatively long delays for our experiment compared with previous single-trial delay detection tasks (Leube et al., 2003; Limanowski, 2017; Limanowski et al., 2020), because our pilot experiments indicated that delay perception was more difficult with our task setup, likely because it required simultaneously maintaining a continuous synchronization of the movements with the pacing rhythm. Each delay level was presented three times, in randomized order, resulting in 24 trials in total. After each trial, we asked our participants to indicate how difficult they felt it was to follow the auditory cue, in comparison to the synchronous practice trial, on a 5-point Likert scale from “very easy” to “very difficult”.

In the second, main part of the experiment, participants also executed continuous, paced grasping movements – now with an added delay to the virtual hand movements that changed repeatedly, following a roving oddball paradigm (Fig. 1D). Participants put their left index finger on the space bar of a computer keyboard, and were instructed to press it as fast as possible every time they detected a change in the visual movement delay (i.e., a change in the mapping of the virtual hand movements to their actually executed movements). Participants were specifically instructed and trained to maintain a stable grasping rhythm with their real hand; i.e., to not alter their movements in response to the delays or delay changes.

During the experiment, each of 8 “baseline” delay levels (ranging from 0ms i.e. synchronous to 700 ms in steps of 100 ms, see above) was presented for 3 to 6 movement cycles (i.e., 6- 12 s), before changing to a larger or smaller delay. The delays changed between ±100-400 ms (in steps of 100 ms), whereby the changes were introduced gradually; i.e., over the course of 0.5 s to prevent sudden jumps in the visual movement. The delays were presented in runs of 147 movements (4.9 mins), each of which contained a predefined, pseudorandomized sequence of 32 delay changes. Thus, we ensured that every run contained 8 delays with each length (3, 4, 5, and 6 movement cycles); and that, within each run, each baseline delay level was presented once in combination with each *absolute* delay change (i.e., ±100-400 ms; 8 delays x 4 changes = 32 delay changes). We focused our analysis on the absolute magnitude of delay changes (±100-400 ms) because we could not systematically compare *relative* i.e., directional delay changes (in- vs decreases) at all levels. In other words, due to the nature of the design, it was not possible to include all combinations of baseline delay and delay changes in- *and* decreases. For instance, at 700 ms, the delay could only decrease, not increase (i.e., there was no change of 700+100 ms, only 700-100 ms). As a control analysis, we tested for potential differences between delay in- vs decreases with a repeated measures ANOVA with the factors delay change (100-400) and direction of change (in- vs decrease) on the detection scores; which showed that detection was not significantly different between delay in- vs decreases (F(3,108)=0.09, *p*=.77). An example of a roving oddball delay sequence can be seen in Figure 1D. Overall, eight such sequences were generated, and presented to each participant in randomized order; resulting in 256 delay changes per participant in total. After each run, participants could take a break until they felt ready for the next one.

### 2.4 Data analysis

To analyze delay change detection performance, we defined a correct delay detection (“hit”) as a button press within two movement cycles (4 s) of a delay change. We chose to include the second movement cycle, because each delay change was gradually introduced (see above), leaving only a partial movement cycle for “detection” otherwise. For consistency, we also tested whether our results could be replicated with detection windows of one and three movement cycles. Button presses outside of this window were categorized as false alarms. If multiple button presses were recorded within 250 ms of a hit or a false alarm (which we (Jain et al., 2015), they were counted as a single button press.

To quantify the overall delay change detection accuracy, we first calculated the *False Alarm Ratio* as the number of false alarms per number of total alarms (i.e., button presses) (Barnes et al., 2007). Note that this is different from the often used False Alarm *Rate* (also called False Positive Rate), which is defined as the number of false alarms per total number of “non-events”. In our case, we could not clearly define “non-events”, as this would apply to any time point not corresponding to a delay change, i.e., to most of the actual movement period.

Then, we calculated a “Detection Score” as a more conservative index of detection performance, punishing the percentage of hits by the False Alarm Ratio. Thus, participants who were more hesitant and thus more accurate, but had lower hits overall, would still receive an adequate score as compared to participants that hit the button at a near random rate. We calculated the detection score as follows:

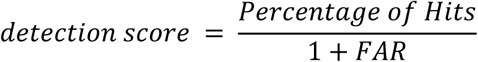

Detection scores were calculated for each of the 32 delay changes (see above). We then inserted the detection scores into a 4 x 8 repeated measures ANOVA, with the factors *Delay Change* (100, 200, 300, 400 ms) and *Baseline Delay* (0, 100, 200, 300, 400, 500, 600, 700, 800 ms). To test for non-normal properties, we used the Shapiro test. Since our data showed non-normal properties, we used an aligned ranks transformation ANOVA for our analysis. The degrees of freedom were calculated using the Kenward-Roger approximation. We also tested for outliers, defined as values of more than three times the interquartile range above the third quartile or below the first quartile; no such extreme values were found. Furthermore, we tested for sphericity using Mauchly’s Test. Since sphericity may be violated for delays, we used Greenhouse-Geisser corrections for our results. When determining the significance of our results, we corrected for multiple comparisons using the Holm-Bonferroni method. Post-hoc, we used a linear regression model (method of least squares) to further evaluate the main effects; i.e., in terms of consistency and directionality.

As an index of tracking performance, we calculated the auditory cue tracking error as the averaged absolute phase shift in time between the normalized mean movement of the participant and the target movement. To test whether tracking performance (i.e., how well participants were able to maintain an accurate and stable grasping rhythm) was influenced by detection performance, we calculated Spearman’s rank correlation coefficients for tracking errors vs the detection scores and vs the reaction times, respectively, averaged over delay levels.

## 3. Results

A repeated measures aligned ranks transformation ANOVA on the difficulty ratings given in the first part of the experiment (i.e., the reported difficulty of tracking the auditory target phase with the grasping movements) showed no significant effect of delay on tracking difficulty (F(7,885)=0.64, p=.72, for the average difficulty ratings of each delay level see Table 1). The difficulty ratings for all delay levels were unaffected by order effects (paired t-tests on ratings given in the first vs last third of the rating blocks, all *n.s.*). Together, this means that participants found the paced movement task itself comparably difficult across all delay levels.

**Table 1.** Means and standard deviations for difficulty ratings, tracking error, detection score and reaction times, for all delay levels.

|  | Delay level |  |  |  |  |  |  |  |
| --- | --- | --- | --- | --- | --- | --- | --- | --- |
|  | 0 ms | 100 ms | 200 ms | 300 ms | 400 ms | 500 ms | 600 ms | 700 ms |
| Difficulty rating [1-5, a.u.] | 2.58<br>±1.15 | 2.51<br>±1.20 | 2.44<br>±1.08 | 2.51<br>±1.17 | 2.53<br>±1.01 | 2.33<br>±1.04 | 2.50<br>±1.17 | 2.59<br>±1.12 |
| Tracking error [s] | 0.23<br>±0.17 | 0.22<br>±0.18 | 0.24<br>±0.18 | 0.23<br>±0.18 | 0.23<br>±0.18 | 0.27<br>±0.18 | 0.27<br>±0.18 | 0.24<br>±0.18 |
| Detection Score [a.u.] | 0.08<br>±0.04 | 0.1<br>±0.03 | 0.12<br>±0.04 | 0.12<br>±0.06 | 0.17<br>±0.08 | 0.14<br>±0.02 | 0.16<br>±0.05 | 0.17<br>±0.04 |
| Reaction Time [s] | 2.14<br>±0.18 | 2.04<br>±0.19 | 2.11<br>±0.27 | 2.06<br>±0.20 | 2.02<br>±0.14 | 1.97<br>±0.30 | 1.82<br>±0.20 | 1.85<br>±0.30 |

In the main experiment, participants were able to maintain regular grasping and tracked the auditory cue well and comparably across delay levels (Fig. 2). On average, they slightly lead the phase of the auditory cue, with an average absolute tracking error of 0.24±0.15 s. Importantly, however, tracking error did not significantly differ between delay levels (F(7,295)=0.23, p=.98, see Table 1). Together, these results suggest that our participants were well able to comply with the task instructions, and that the amount of delay of the virtual hand itself did not significantly influence tracking (grasping) performance, nor the perceived task difficulty.

**Figure 2.**
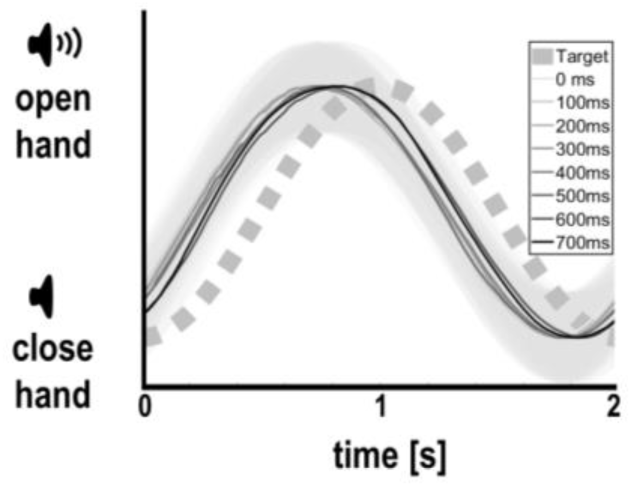
Participants’ average hand movements per delay level (with standard deviations in gray shading), relative to the instructed rhythm prescribed by the auditory cue. Participants tracked the auditory target rhythm comparably well across all delay levels, albeit overall leading slightly (see Results for details). Thus, visual delay per se did not bias grasping or tracking performance.

To validate the overall delay change detection accuracy of our participants, we first tested it against the chance level. The average False Alarm Ratio of our participants was 0.45±0.10, which was significantly lower than the ratio that would have resulted from random button pressing (i.e., >0.67; t(36)=13.13, p<.001). This suggests that, overall, the participants were able to notice the delay changes and pressed the button accordingly.

An aligned ranks transformation ANOVA on the detection scores showed a significant main effect of Delay Change; i.e., the magnitude of the change in the delay of the visual hand movement (F(3,1116)=19.48, p<.001, ƞ^2^=.05). A post-hoc linear regression showed that these effects could be linearly approximated (Fig.3A, R²=.07, Slope= 2.58·10^-4^, df= 35, p<.05; for average detection scores see Table 2). However, there was no significant effect of Delay Change on the corresponding reaction times (Fig. 3B, for means and standard deviations see table 2), while the interaction effect was significant (F(21,1116)= 3.505, p<.001, ƞ^2^=.06). In sum, as expected, larger delay changes were consistently detected better (albeit not significantly faster) than smaller ones.

**Figure 3.**
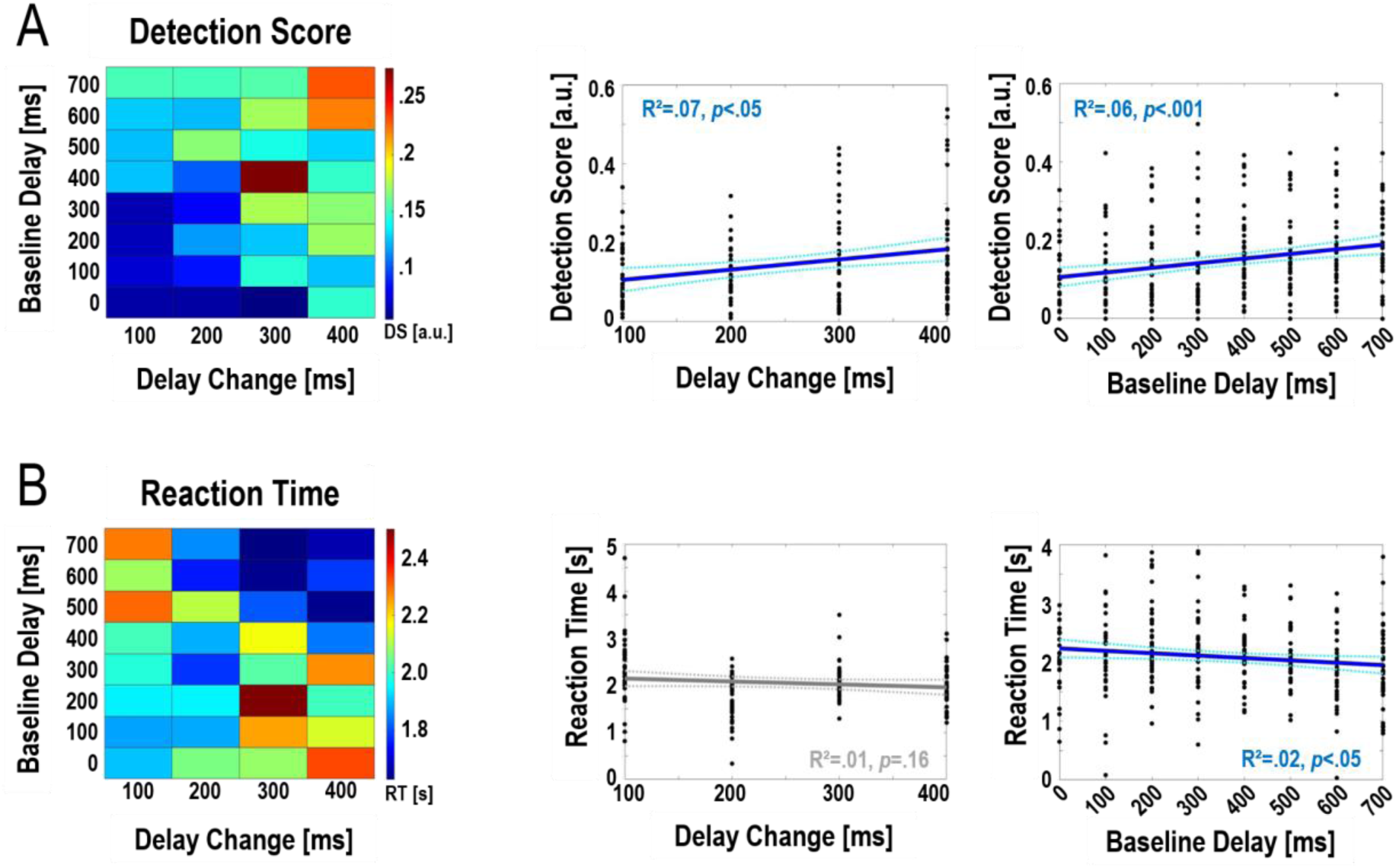
Accuracy and speed of detecting changes in visual movement feedback delay during continuous grasping. The left plots show the average detection scores (**A**) and reaction times (**B**) for each of the 32 combinations of delay change magnitude and baseline delay. **Detection accuracy as quantified by the Detection Score was significantly affected by the amount of delay present during movement (main effect Baseline Delay, *p*<0.001) and the magnitude of the change in delay (main effect Delay Change, *p*<0.001). Reaction times were significantly affected by Baseline Delay (main effect, *p*<0.001). The right plots show post-hoc linear regression analyses (see Results for details).** Overall, these analyses showed that larger delay changes were detected significantly more accurately (and slightly but not significantly faster) than smaller changes. Furthermore, change detection was significantly better and faster when participants were moving at larger, compared to smaller, delays.

**Table 2.** Means and standard deviations for detection scores and reaction times, for the four different absolute magnitudes of delay change, respectively.

|  | Delay Change (absolute) |  |  |  |
| --- | --- | --- | --- | --- |
|  | 100 ms | 200 ms | 300 ms | 400 ms |
| Detection Score [a.u.] | 0.10 ±0.03 | 0.11 ±0.03 | 0.15 ±0.06 | 0.16 ±0.04 |
| Reaction Time [s] | 2.07 ±1.23 | 1.81 ±0.84 | 2.04 ±0.76 | 2.01 ±0.77 |

Secondly, there was a significant main effect of Baseline Delay on detection scores (F(7,1116)= 11.76, p<.001, ƞ^2^=.07) and reaction times (F(7,1116)=6.56, p<.001, ƞ^2^=.04); as well as a significant interaction between Baseline Delay and Delay Change (F(21,1116)=2.09, p<.05, ƞ^2^=.04 Fig.3B). Post-hoc linear regression analyses showed, again, that these effects could be linearly approximated (detection scores, Fig.3A: R²=.06, Slope= 1.18·10^-4^, df= 35, p<.001; reaction times, Fig.3B: R²=.02, Slope= -4.05·10^-4^, df= 35, p<.05, for means see Table 1. In sum, delay change detection was overall better and faster at larger > smaller delays; i.e., the more the visual movement feedback was delayed during movement when the change occurred.

The above significant main and interaction effects on detection performance were also significant when limiting the analysis to a detection window of only the first movement cycle (all Fs>2.2, all ps<.01) or the first three movement cycles (all Fs>4.3, all ps<.001). The significant main effect of Baseline Delay on the reaction times, and the interaction effect, were also replicated when limiting the analysis to the first movement cycle (all Fs>3.54, all ps<.001) or the first three movement cycles (all Fs>1.55, albeit not reaching statistical significance, ps<.06). The slopes of the respective linear regressions confirmed the directionality of the effects as in our main analysis. In short, the width of our detection window did not bias our results.

Finally, we tested whether the participants’ average detection performance covaried with how well they were able to maintain stable tracking. With this, we aimed to detect a possible relationship between delay detection and tracking error despite the fact that tracking performance on average was comparable across delay levels (cf. Fig. 2). This analysis revealed a significant negative correlation between detection accuracy and tracking performance: Participants who tracked the target rhythm overall better (i.e., had a smaller average tracking error) detected less of the delay changes (ρ=.40, p<.05, Fig. 4). The correlation between reaction times and tracking error showed a similar directionality, but was not significant (smaller tracking error ∼ longer reaction times to delay changes; ρ=-.05, p=.78).

**Figure 4.**
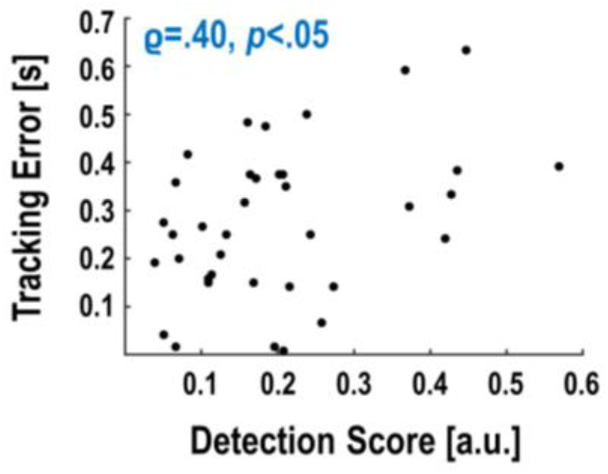
Spearman correlation between detection score and tracking error of the individual participants. **Participants with better detection scores had a larger tracking error; i.e., they followed the auditory target cue worse with their grasps.**

## 4. Discussion

Using a virtual reality based continuous hand movement task, we tested whether the detection of changes in visuomotor delay depended on the amount of delay currently present during movement. As expected, we found that larger delay changes were detected significantly better than smaller ones. Somewhat surprisingly, however, delay changes were detected significantly better and faster when participants moved under larger > smaller delays. In other words, our results suggest a higher sensitivity to changes in visual movement feedback delay when there already is a substantial amount of delay present during movement. This seems to indicate that the Weber-Fechner Laws do not apply to the perception of (temporal) visuomotor mapping; as they would predict the opposite; i.e., better sensitivity to delay changes under smaller delays.

Results inconsistent with the Weber-Fechner Laws have been reported in other domains before, where they have been interpreted as indicating, for instance, saturation effects (Augustin, 2008; Augustin & TANJA, 2008; Carriot et al., 2021; Doble et al., 2003). In our case, we propose two complementary explanations for our results:

Firstly, it is well established that increasing visuomotor delay reliably reduces the self-attribution of the observed movements (also called a loss of control or “agency” over those movements (Krugwasser et al., 2019; Wen & Imamizu, 2022) and the subjective embodiment of the seen moving body part (also called a loss of “ownership”; see (Dummer et al., 2009; Farrer et al., 2008; Leube et al., 2003; Tsakiris et al., 2006). Thus, it is reasonable to assume that, in our experiment, participants experienced a relatively stronger subjective embodiment of the virtual hand under small delays. After our experiment, seven of our participants spontaneously reported having perceived up to half of all virtual movements as synchronous to their own movement (i.e., much more than actually were synchronous) – several of those participants reported a feeling of ownership over the virtual hand in those synchronously perceived periods. While these were only spontaneous post-hoc reports, they tentatively support the idea of embodiment biasing delay detection in our task.

Experienced embodiment can substantially affect attentional control and task performance in virtual reality based movement tasks, as shown e.g. by Iwasaki and colleagues (2022). In our case, one such effect could be intersensory conflict attenuation. As demonstrated in the ‘rubber hand illusion’ (Botvinick & Cohen, 1998; Limanowski, 2022), the embodiment of an alternative (fake or virtual) limb biases the subjective perception of visuoproprioceptive mismatches between real and fake limbs, attenuating intersensory conflict through updates to the corresponding neuronal representations (Limanowski, 2022). Since visuomotor delays are, effectively, also visuoproprioceptive mismatches, we speculate that a similar intersensory conflict attenuation could occur at smaller, compared with larger delays. In other words, an increased embodiment of the virtual hand at smaller delay levels could have increased the tolerance for visuomotor mismatches, and thus, impaired delay detection performance. An increased embodiment of the virtual hand under small > large delays could also have distorted the perception of movement timing more generally (Desantis et al., 2016; Haering & Kiesel, 2015; Rohde & Ernst, 2016).

A complementary influence on detection performance could have resulted from larger visuomotor delays being more salient stimuli *per se*. This saliency effect could stem from the fact that these delays constitute ‘non-standard’ visuomotor mappings; i.e., mappings that violate the life-long learned associations (of congruence) between motor commands and visual movement feedback (Quirmbach & Limanowski, 2024; Yon et al., 2018). Thus, larger delays could have captured more attention—onto the visuomotor relationship, or more generally onto the visual movement feedback—than smaller delays. This would likely have improved delay detection performance at larger delay levels. Note that this interpretation complements, rather than contradicting, the idea of an embodiment-related increased tolerance for changes at lower delay levels. However, as we did not explicitly assess subjective embodiment or attentional allocation, both of the above interpretations have to be explicitly tested in future experiments.

Finally, note that our task design implied maintaining focus on the real hand movements, to track the auditory rhythm, while the virtual hand moved incongruently. Visual body movements that are incongruent with one’s actual movements and movement goals can capture attention and lead to behavioral deficits—this is sometimes also referred to as visuomotor interference (Blakemore & Frith, 2005; Brass et al., 2001; Kilner et al., 2003; Limanowski & Friston, 2020; Metral & Guerraz, 2019). The distracting effect of incongruent visual stimuli may be particularly strong when they depict one’s own, delayed movements (Salomon et al., 2013). The positive correlation between detection performance and tracking error suggests that participants who used more cognitive-attentional resources to maintain stable tracking lacked those resources for the detection task. The ‘dual task’ nature of our design could also explain the relatively low overall detection performance we observed, in contrast to other studies using single-movement designs (Farrer et al., 2008; Leube et al., 2003). Interestingly, tracking performance did not differ between delay levels. This could suggest that our participants resisted or counter-acted any potential biasing effect of visual feedback delay on movement execution (cf. Limanowski & Friston, 2020b). An interesting question (which we, however, cannot answer) is whether an increased embodiment of the virtual hand would lead to stronger visuomotor interference effects (hence expected at smaller delays); or whether larger delays *per se* elicit stronger visuomotor interference effects.

Our study raised further questions that should be followed up by future work. Due to the nature of our design, we could not systematically compare detection and reaction times to delay in- vs decreases. Although in a control analysis, we did not find any significant differences in detection scores between delay in- vs decreases (see Methods), this needs to be addressed in detail by future work. Furthermore, it has been shown that, when moving under visuomotor delay, visual movement feedback is processed differently depending on whether vision is currently task-relevant or a distractor (Limanowski et al., 2020; Limanowski & Friston, 2020). An open question is whether visuomotor delay *changes* are processed differently depending on whether they are task-relevant or -irrelevant. Furthermore, we observed substantial performance differences (in tracking and detection) between participants; future work should look into the possible causes for such inter- but also intrapersonal variability. For instance, one factor influencing overall performance could have been prior experience with VR scenarios, which we did not assess. Visuomotor temporal incongruence can in principle also be implemented by leading, not lagging, vision; in the context of self-initiated hand movements, however, this is associated with a number of problems related to the predictability of the visual trajectories before actual movement. Finally, visual feedback delay is only one kind of visuomotor incongruence; future work should test whether other manipulations, such as spatial offset or gain scaling, produce similar results.

To conclude, we have shown that, during bodily action in virtual reality, the current degree of visuomotor delay can bias the perception of changes in the magnitude of this delay. This bias does not seem to follow the Weber-Fechner-Laws. Instead, bodily action under relatively small delays may entail some degree of tolerance for delay changes resulting from intersensory conflict attenuation due to a stronger embodiment of the (relatively more congruently moving) virtual hand; whereas large delays may capture (visual) attention due to their violation of visuomotor predictions. Thus, our results highlight the importance of cognitive-attentional factors in visuomotor processing; and the need to consider these factors when designing cyber-physical interactions with variable delays.

## Declarations

### Ethics approval

The experiment was approved by the ethics committee of the Technische Universität Dresden and conducted in accordance with this approval (SR-EK-174032021).

### Consent to participate

Voluntary informed consent was obtained from all participants.

### Consent for publication

The authors affirm that participants provided informed consent for publication of anonymized data.

### Availability of data and materials

Data or materials for the experiments reported here are available upon request. None of the experiments was preregistered.

### Code availability

Code for this experiment is available upon request.

## Acknowledgments

This work was funded by the German Research Foundation (DFG, Deutsche Forschungsgemeinschaft) as part of Germany’s Excellence Strategy – EXC 2050/1 – Project ID 390696704 – Cluster of Excellence “Centre for Tactile Internet with Human-in-the- Loop” (CeTI) of Technische Universität Dresden. JL was supported by a Freigeist Fellowship of the VolkswagenStiftung (AZ 97-932).

